# A central control circuit for encoding perceived food value

**DOI:** 10.1101/368969

**Authors:** Michael Crossley, Kevin Staras, György Kemenes

## Abstract

Hunger state can substantially alter the perceived value of a stimulus, even to the extent that the same sensory cue can trigger antagonistic behaviors. How the nervous system uses such graded perceptual shifts to select between opposed motor patterns remains enigmatic. Here we challenged food-deprived and satiated *Lymnaea* to choose between two mutually exclusive behaviors, ingestion or egestion, produced by the same feeding central pattern generator. Decoding the underlying neural circuit reveals that the activity of central dopaminergic interneurons defines hunger state and drives network reconfiguration, biasing satiated animals towards the rejection of stimuli deemed palatable by food-deprived ones. By blocking the action of these neurons, satiated animals can be reconfigured to exhibit a hungry animal phenotype. This centralized mechanism occurs in the complete absence of sensory retuning and generalizes across different sensory modalities, allowing food-deprived animals to increase their perception of food value in a stimulus-independent manner to maximize potential calorific intake.

## Introduction

Hunger is a potent regulator of animal behavior. Periods of food-deprivation can increase risky decision making, for example ignoring potential threats in the environment (Burnett et al., 2016; Gaudry and Kristan, 2009; Ghosh et al., 2016; Jikomes et al., 2016; Padilla et al., 2016; Schadegg and Herberholz, 2017) or ingesting potentially harmful food (Gillette et al., 2000; Inagaki et al., 2014; LeDue et al., 2016). In the latter example, animals must make dynamic decisions about the cost-benefit of food intake relative to their motivational state, and select an appropriate behavior; in some cases, the complete reversal of the motor pattern, from ingestion to rejection. How the nervous system encodes internal state, makes decisions about the perceived value of a stimulus and selects between alternative behavioral responses by motor network reconfiguration, are significant questions in neurobiology.

Previous studies have demonstrated that hunger state can modulate behavioral responsiveness to a stimulus by influencing sensory pathways (Inagaki et al., 2012; Inagaki et al., 2014; Ko et al., 2015; LeDue et al., 2016; Root et al., 2011). Although this provides an efficient tuning mechanism for generating simple graded changes in a particular behavior, whether it could underlie more complex decisions resulting in complete behavioral reversal is unknown. A second possibility, raised by the observation that in both vertebrates and invertebrates, ingestion and egestion are driven by a single CPG (Jing and Weiss, 2002; Li et al., 2016), is that the control switch is central not peripheral. In this case motivational state would act on the pattern-generator or a connected higher-order network directly, to select the appropriate behavior. This type of central control switching mechanism would endow the system with the ability to generalize behavioral selection to multiple stimuli without the need to retune all the sensory pathways that mediate them.

Here we examine these possibilities using *Lymnaea*, one of the best-characterized models for studying feeding control and its neural mechanisms (Benjamin, 2012; Harris et al., 2012; Kemenes et al., 2001; Marra et al., 2013; Pirger et al., 2014; Staras et al., 1998). This animal generates clearly definable ingestion and egestion behaviors using a well-identified CPG (Straub et al., 2002; Vavoulis et al., 2007) that drives a highly complex feeding musculature consisting of 46 muscles (Benjamin, 2012). This system offers a unique entry point for elucidating fundamental mechanisms that encode changes in motivational state and direct the expression of relevant animal behavior (Crossley et al., 2016; Dyakonova et al., 2015; Staras et al., 2003).

A forced choice paradigm was employed to demonstrate that a single potential food stimulus drives mutually exclusive behaviors, either ingestion or egestion, that depends on the hunger state of the animal. Using *in vivo* and *in vitro* approaches we show that this behavioral selection utilizes a higher-order switch mechanism acting directly on the feeding network to drive a motivational state dependent choice, without sensory retuning. This is achieved by activity in hunger-state encoding dopaminergic interneurons, which reconfigure the feeding network to bias satiated animals towards the rejection of stimuli deemed palatable by food-deprived ones. We also show that this higher-order control allows generalization across different input modalities. In this way, according to its hunger state, an animal can universally adjust the level of risk it is prepared to accept when deciding whether to consume ambiguous material that has food-like cues. Our study reveals a central neuronal switch mechanism that endows the animal with the advantageous ability to make adaptive choices regarding food intake within its highly unpredictable sensory habitat.

## Results

### Circuit level expression of mutually exclusive feeding behaviors

*Lymnaea* can perform two types of feeding behavior, ingestion and egestion, which serve opposing functions. Both behaviors utilize the same feeding structures (mouth, buccal mass and radula) (Fig. 2a, b) and are therefore mutually exclusive. Application of an appetitive stimulus to the lips (lettuce or sucrose) elicits ingestion bites (Fig. 1a, b, top panels), whereas application of an aversive stimulus (pinch to the esophagus) triggers egestion where the movement of key feeding structures is reversed (Fig. 1b, bottom panels).

**Fig. 1.**
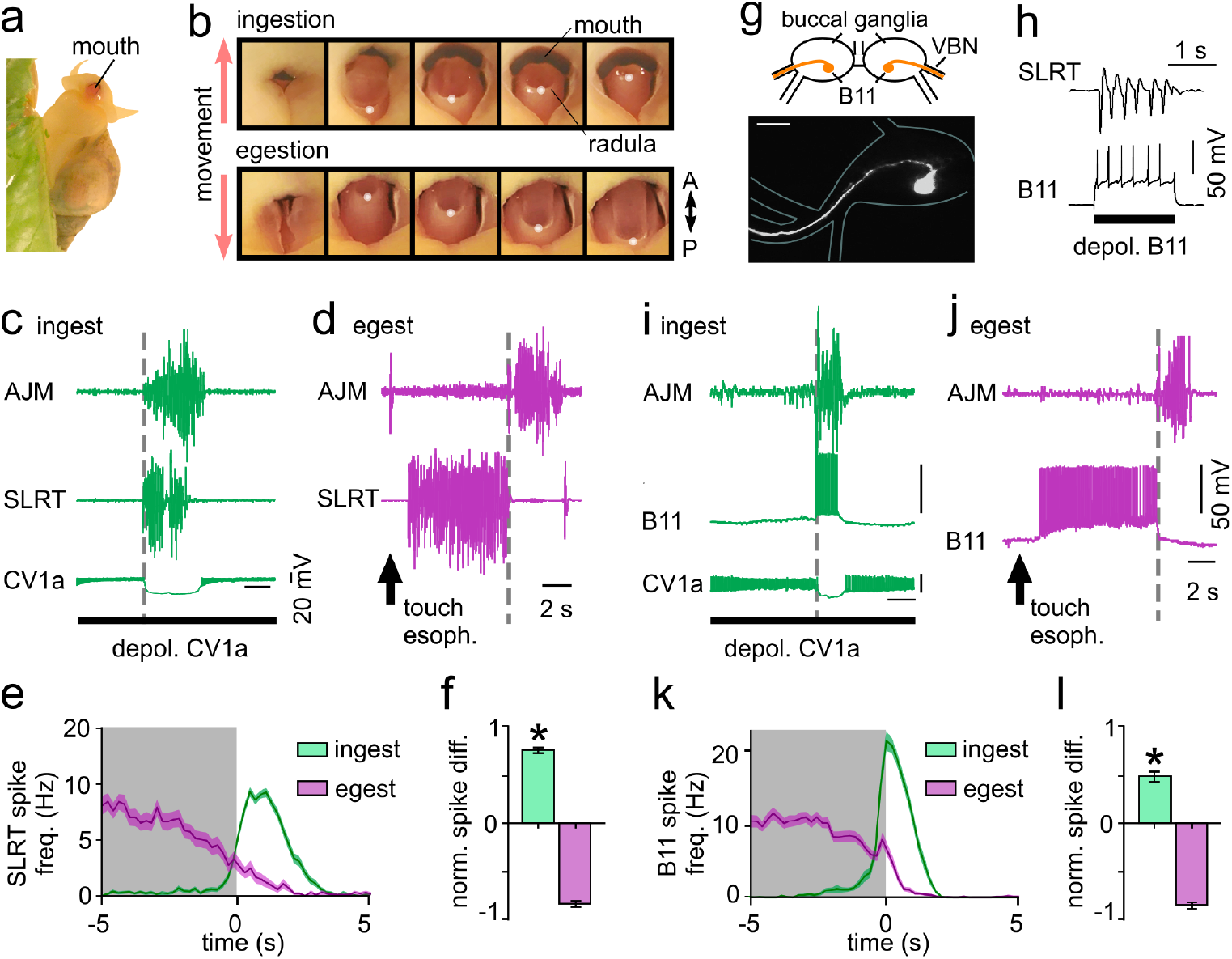
*In vitro* correlates of ingestion and egestion behavior. (**a**) *Lymnaea* biting at water surface. (**b**) Mouth movements during ingestion or egestion. White dot indicates the distal tip of the radula tracked in analysis. (**c-d**) AJM and SLRT activity during ingestion (c) and egestion (d) cycle. Gray dotted line represents the onset of burst of activity on AJM. (**e**) Average frequency of SLRT activity during ingestion (green) (n = 105 cycles, 10 preps) and egestion (purple) (n = 51 cycles, 8 preps). Line and shading show mean ± SEM. (**f**) Normalized difference score of SLRT activity after vs before AJM activity onset with significant difference between ingestion (0.77 ± 0.05) and egestion (−0.83 ± 0.05, unpaired t-test, p < 0.0001). Data shows mean ± SEM. (**g**) Morphology and schematic of B11 motoneuron in the buccal ganglia showing single projection leaving ganglion via ventral buccal nerve (VBN) towards buccal mass and SLRT muscle. Morphology confirmed in n = 3 cells. Scale bar 100 μM. (**h**) Spikes artificially triggered in B11 cause 1:1 activity on SLRT muscle. (**i-j**) B11 and AJM activity during ingestion (i) and egestion cycles (j). (**k**) Average B11 spike frequency during ingestion (green) (n = 69 cycles, 9 preps) and egestion (purple) (n = 78 cycles, 10 preps) cycles. Line and shading are mean ± SEM. (**l**) Normalized difference scores of B11 activity after vs before AJM activity onset showed a significant difference between ingestion (0.49 ± 0.05) and egestion (−0.87 ± 0.03, unpaired t-test, p < 0.0001). Data shows mean ±SEM.

**Fig. 2.**
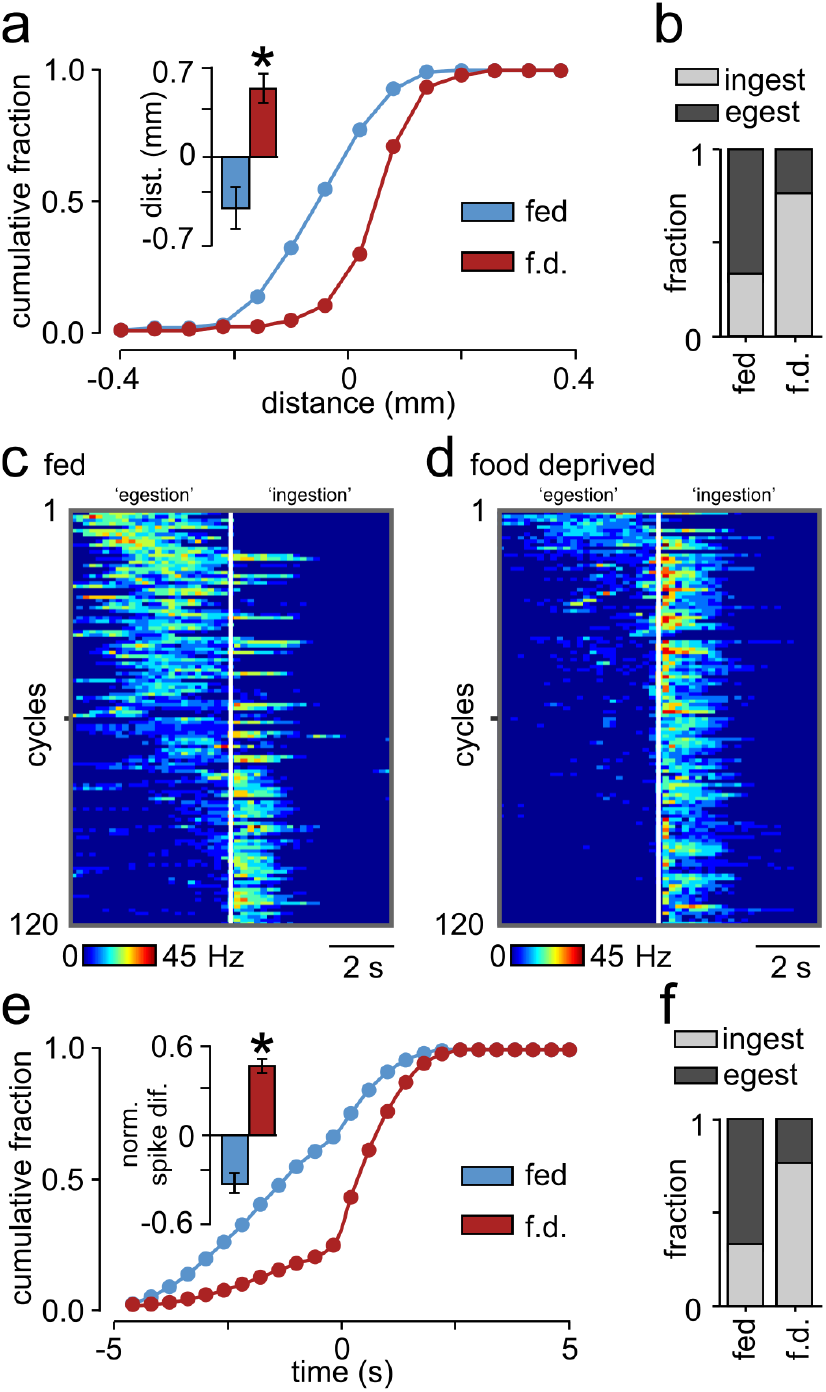
Choice between ingestion and egestion depends on hunger-state. (**a**) Cumulative frequency plot of radula movements in response to tactile probe from fed and food-deprived animals. Bar chart of behavioral response to tactile probe shows a significant difference in radula movements between fed (−0.41 ± 0.17 mm, n = 18) and food-deprived (0.55, ± 0.12 mm, n = 21) animals (Mann Whitney test, p < 0.0001). Data shows mean ± SEM. (**b**) Comparison of fraction of ingestion and egestion bites in response to tactile stimulus with significant difference between fed and food-deprived responses (Fisher’s exact test, p < 0.0001). (**c-d**) Heat plots of B11 activity in multiple trials during *in vitro* cycles from fed (c) and food-deprived (d) preparations. White line represents onset of AJM burst. Data ordered from high to low activity prior to AJM onset. (**e**) Cumulative frequency plot of B11 activity during cycles from fed and food-deprived preparations. Bar chart showing normalized B11 spike frequency before vs after AJM burst onset (n = 120 cycles from 12 preparations for both conditions) showing a significant difference between fed (−0.33 ± 0.12) and food-deprived (0.47 ± 0.09) cycles (unpaired t-test, p < 0.0001). (**f**) Comparison of the fraction of ingestion and egestion cycles. There was a significant difference between fed and food-deprived preparations (Fisher’s exact test, p < 0.0001).

To confirm that this opposed behavioral expression was the consequence of a reversal of the motor control patterns *in vitro*, we carried out experiments in which we co-recorded from two key radula muscles, the anterior jugalis muscle (AJM) and the supralateral radula tensor muscle (SLRT)(Carriker, 1946; Rose and Benjamin, 1979) in a reduced preparation. Cycles evoked by the command-like interneuron CV1a, an established part of the food-signaling pathway (Kemenes et al., 2001; McCrohan, 1984) and see Supplementary Figure 1a, b), or by appetitive stimuli applied to the lips, produced a single burst of synchronized activity per cycle in both muscles (Fig. 1c, e, f and Supplementary 1c). By contrast, egestion behavior, evoked by tactile stimulation of the esophagus, an aversive stimulus (Supplementary Figure 1b) produced highly asynchronous muscle activity, with SLRT activation preceding AJM (Fig. 1d-f). As such, differential AJM/SLRT activity expression provides a robust *in vitro* correlate of ingestion/egestion. Next, we looked for neurons in the CNS that could underlie these opposing muscle activity patterns. Our search identified a new type of motoneuron, B11 (Fig. 1g, h), which both projected to SLRT and robustly activated it in a 1:1 manner. Furthermore, B11 activity mirrored SLRT activity in both ingestion and egestion cycles (Fig. 1i-l) showing in-phase activity with the AJM during ingestion cycles but out-of-phase activity during esophageal-activated egestion cycles. As such, B11 is differentially recruited in the two different behaviors and its activity provides an important central readout of ingestion/egestion expression *in vitro*.

### Choice between ingestion and egestion depends on hunger-state

Two opposed behaviors with known neural correlates provide a powerful system to examine the effects of hunger-state on behavioral selection. To investigate this, we devised a type of forced-choice paradigm where animals in different motivational states are made to choose between ingestion and egestion behaviors based on the presence of a single type of stimulus. Animals were in one of two states, either fed (food available *ad libitum*) or food-deprived (4 days starved), and were tested for their behavioral response to a tactile stimulus with food-like properties (Crossley et al., 2016). In both groups, this stimulus was presented to the mouth as the animal performed an ingestion bite. The radula movement that followed was then classified as either ingestive or egestive, providing a readout of the animal’s judgement regarding the potential value of the stimulus. In fed animals, the tactile stimulus evoked net egestive responses while predominantly ingestive responses were recorded in food-deprived animals (Fig. 2a, b). Thus, a single stimulus can elicit one of two mutually exclusive behaviors depending on hunger-state: fed animals perceive the stimulus as inedible while their food-deprived counterparts initiate ingestive feeding responses.

Next, we tested whether food-deprivation is associated with a generalized shift in the state of the feeding network resulting in a switch from egestion to ingestion. As a readout of network state we used the occasional spontaneous cycles recorded *in vitro* and classified them as ingestion or egestion patterns based on B11 versus AJM activity (Fig. 2c, d). Notably, we found that preparations from fed animals were highly biased towards egestion activity, while hungry animals were associated with ingestive patterns (Fig. 2e, f). We reasoned that this might be explained by an inability of hungry animal preparations to generate egestion cycles at all. To test this, we recorded their B11 responses to esophageal stimulation (Supplementary Figure 2), and demonstrated that egestive responses occurred at the same level as observed in fed animals (food-deprived, n = 8, −0.8 ± 0.06, fed, n = 10, −0.87 ± 0.03, normalized difference scores, see Fig. 1 legend for detail of measure, unpaired t-test p > 0.05). Thus, during periods of food-deprivation, there is a shift in the state of the feeding network from egestion towards ingestion, presumably associated with a motivational state-dependent change in the central circuits driving these two opposing behaviours.

### Higher-order interneurons encode hunger-state and select motor patterns

Next, we carried out an extensive characterization of the CNS to identify elements that might drive this hunger-state dependent decision-making circuit. In particular, our search identified a pair of buccal to cerebral interneurons (Fig. 3a) with key characteristics consistent with this function and therefore termed Pattern Reversing Neurons (PRNs). First, activating the egestion pathway *in vitro* elicited a strong burst of spikes in a PRN, which coincided with B11 activity prior to AJM activity (Fig. 3b). Moreover, at the onset of the retraction phase, activity in PRN and B11 both ceased. Second, during ingestion cycles, PRN showed no spiking activity (Supplementary Figure 3a) indicating that it was selectively active during egestion cycles. Third, artificial activation of a single PRN was sufficient to drive exclusively egestive motor programs (Fig. 3c and Supplementary Figure 3b) and there was no significant difference between PRN driven cycles and sensory-triggered egestion (PRN, n = 8, −0.77 ± 0.04, normalized difference score, unpaired t-test p > 0.05). Likewise, CV1a, a driver of ingestive feeding cycles, exhibited no activity during PRN driven cycles (Supplementary Figure 3c). To ascertain whether PRN played a role in the motivational-state dependent biasing of motor-pattern selection, PRN activity was recorded during spontaneous cycles in preparations from fed and food-deprived animals. We found that these cells exhibited significantly higher firing rates per cycle and significantly more activity cycles in total, in fed versus food-deprived preparations (Fig. 3d-f). Taken together, these results demonstrate that a key central control element of the egestion network is in a relative upstate in fed preparations compared with their food-deprived counterparts. Next, we tested whether PRN activity was the source of the bias towards egestion patterns in fed preparations by inhibiting these cells during cycles. We found that hyperpolarizing both PRNs alone was sufficient to switch fed preparations from an egestion to ingestion pattern (Fig. 3g). Notably, preventing somatic spikes in both PRNs did not block esophageal driven egestion (Supplementary Figure 3f) suggesting that while other elements are involved in the basic ability to egest, PRN has a critical hunger-state specific function. Thus, we conclude that these pivotal command-like interneurons encode *in vitro* hunger-state, serving as a central switch mechanism for motor-pattern selection.

**Fig. 3.**
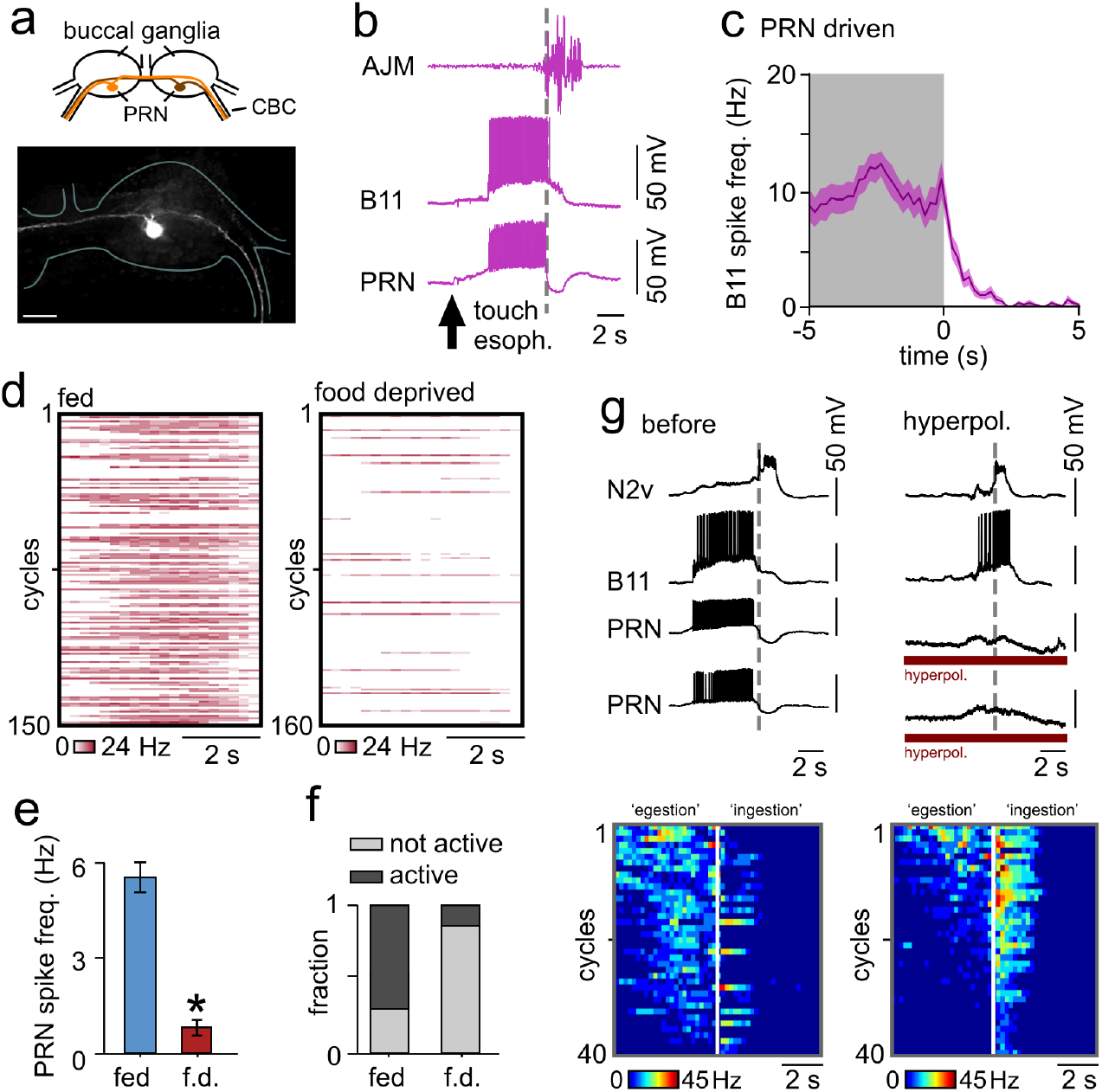
PRN encodes hunger-state and drives network reconfiguration. (**a**) Morphology of Pattern Reversal Neurons (PRNs) with cerebral buccal connective (CBC) projection, confirmed in n = 6 cells. Scale bar 100 μM. (**b**) Trace of PRN activity during esophageal-driven egestion, co-recorded with B11 and AJM. PRN is active in phase with B11 but not AJM. Gray dotted line shows onset of AJM burst. (**c**) Average spike frequency of B11 activity during PRN driven cycles (see Supplementary Figure 3B)(n = 36 cycles, 8 preparations). Line and shading are mean ± SEM. (**d**) Heat plots of PRN firing rates during *in vitro* cycles in fed (left)(n = 150 cycles, 15 preps) and food-deprived (right)(n = 160 cycles, 16 preps) preparations. (**e-f**) Statistical analysis. Fed preparations had more PRN activity per cycle (unpaired t-test, p < 0.0001) and more active cycles (Fisher’s exact test, p < 0.0001) than food-deprived. Data shows mean ± SEM. (**g**) Representative traces of an *in vitro* egestion cycle from fed preparation (left). B11 is only active in protraction phase (see Supplementary Figure 3d, e). Hyperpolarizing PRNs switches B11 activity to ingestion-like pattern (right). Gray dotted lines show retraction phase onset. Heat plots of B11 activity before (left) and during (right) PRN hyperpolarization. White line shows onset of AJM burst. Data ordered from high to low (n = 40 cycles, 4 preps) with significant B11 difference score before vs during PRN hyperpolarization (before: −0.62 ± 0.06, during: 0.4 ± 0.15, paired t-test p = 0.002).

### Switching from a satiated to a hungry phenotype *in vitro* and *in vivo*

To test whether PRNs account for behavioral selection *in vivo*, we first identified the transmitter used by these cells to drive egestion. Previous mapping studies have revealed a single pair of unidentified dopaminergic neurons on the ventral surface of the buccal ganglia (Croll et al., 1999; Elekes et al., 1991; Vaasjo et al., 2018) and we hypothesized that these might be PRNs. Electrophysiology and double-labeling experiments using Alexa Fluor and dopamine antibody staining supported this idea (Fig. 4a). Moreover, experiments showed that the strong PRN→B11 monosynaptic excitatory connection (Fig. 4b, left), which accounts for B11 activation during egestion cycles, was blocked with the D2 receptor antagonist, sulpiride (n = 9)(Fig. 4b right and Supplementary Figure 4a). Thus, we concluded that dopamine is PRN’s transmitter, and next examined whether sulpiride application could mimic the modulation of motor-program selection in fed preparations seen when PRN activity is suppressed (see Fig. 3g). Specifically, we recorded spontaneous cycles on B11 and AJM in fed preparations before and after bath-application of sulpiride to the CNS (Fig. 4c). As expected, egestion cycles predominated in the pre-treatment condition, but following sulpiride application, we observed a striking change in cycle expression, with a switch to almost exclusively ingestive cycles (Fig. 4c-e). Therefore, sulpiride can reconfigure the network state from egestion to ingestion *in vitro*. As before, we demonstrated that this was not explained by an inability of the feeding system to generate egestion cycles at all when sulpiride-treated; tactile stimulation of the esophagus was still able to drive egestion cycles in the presence of the blocker (Supplementary Figure 4b). Taken together, this suggests that sulpiride application *in vitro* inhibits PRN’s state-dependent control pathway and biases the preparation towards the selection of ingestion cycles.

**Fig. 4.**
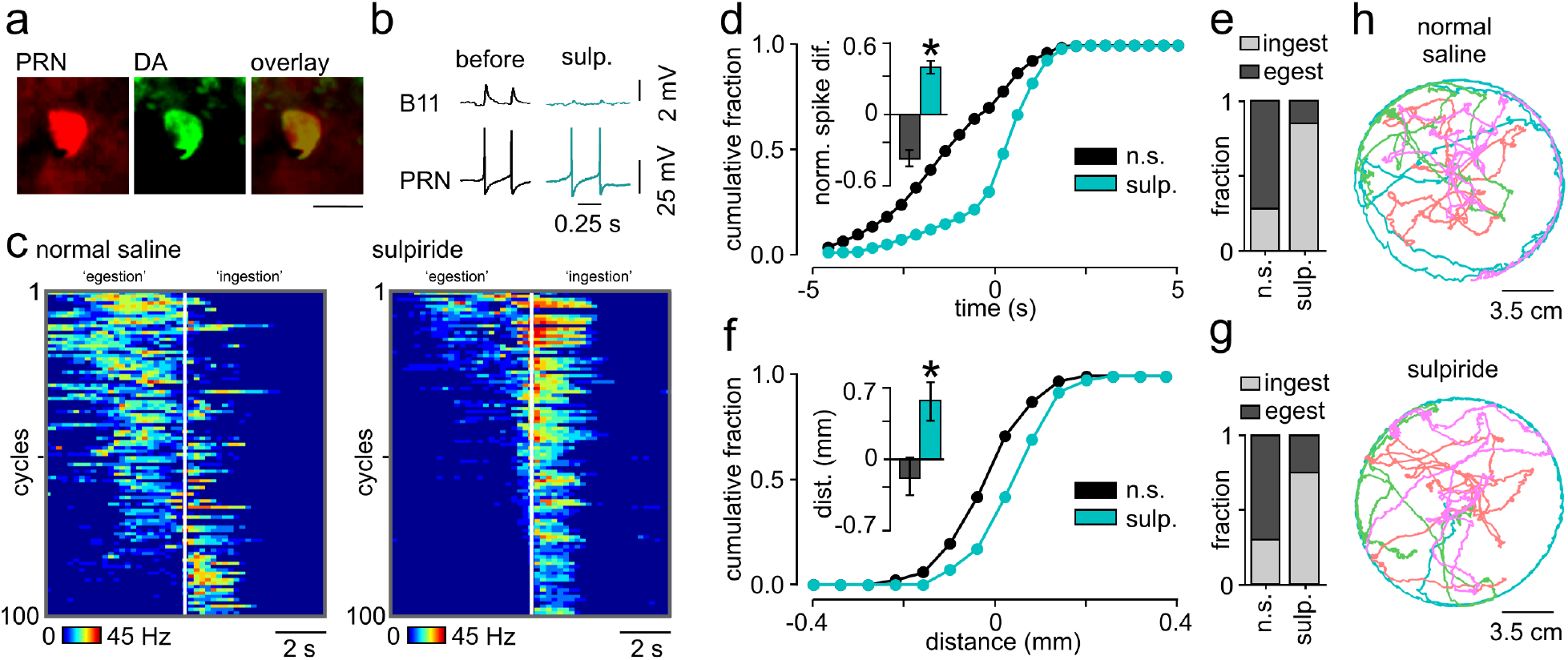
Switching from satiated to hungry phenotype with dopamine block. (**a**) Double labelling of PRN (Alexa Fluor, left) and dopamine neuron (middle) and overlay (right). Confirmed in n = 4 preparations. Scale bar, 25 μM. (**b**) Evoked spikes in PRN caused 1:1 EPSPs on B11 before (left) and after 10^-4^ M sulpiride application (right). (**c**) Heat plot of B11 activity *in vitro* in fed preparations before (left) and after (right) sulpiride application. White line shows onset of AJM burst. Data ordered from high to low activity prior to AJM onset (n = 100 cycles from 10 preparations for both conditions). (**d**) Cumulative frequency plot of B11 activity during cycles in normal saline or sulpiride. Bar chart (mean ± SEM) of normalized B11 spike frequency before vs after AJM activity shows difference between cycles in normal saline (−0.34 ± 0.14) and sulpiride (0.38 ± 0.07, paired t-test, p < 0.05). (**e**) Comparison of fraction of ingestion and egestion cycles for normal saline or sulpiride (Fisher’s exact test, p < 0.0001). (**f**) Cumulative frequency plot of radula movements in response to tactile probe for animals injected with saline (−0.16 ± 0.17 mm, n = 16) and sulpiride (0.51 ± 0.18 mm, n = 16) injected animals (Mann Whitney test, p < 0.05). Data shows mean ± SEM. (**g**) Comparison of the fraction of ingestion and egestion responses to tactile stimulus (normal saline vs sulpiride, Fisher’s exact test, p < 0.0001). (**h**) Representative trajectories of saline (top) and sulpiride-injected (bottom) animals (n = 4) over 30 mins. There was no significant difference in total distance traversed between groups (saline: 117.1 ± 71.74 cm, n = 15, sulpiride: 118.4 ± 65.21 cm, n = 15, unpaired t-test, p = 0.89).

Next, we investigated whether the same pharmacological intervention could determine behavioral selection *in vivo*. Fed animals received the same behavioral test as previously (see Fig. 2) but were injected with either normal saline or sulpiride prior to testing. As expected, in saline-injected animals, the tactile stimulus triggered predominantly egestive responses, but notably, in sulpiride-injected animals, responses were ingestive (Fig. 4f, g). To address the possibility that changes in another parameter of appetitive or consummatory behavior was driving this effect, we characterized the influence of sulpiride on the animal’s food searching locomotion (Fig. 4h), appetitive-bite sampling behavior and consummatory behavior in response to sucrose (Supplementary Figure 4c, d). None of these parameters showed significant differences between treated and control animals. Thus, sulpiride provides a targeted effect on the motivational state-dependent behavioral selection between ingestion and egestion within the feeding network, and effectively switches satiated animals to a hungry animal phenotype. These results show that PRN encodes hunger-state and acts as a central switch for behavioral selection *in vivo*.

### Sensory retuning is not used for behavioral selection

A previously identified mechanism for encoding hunger state-driven adaptive changes in behavior relies on a retuning of the output gain of sensory input pathways (Su and Wang, 2014). To test whether a similar gain control process is also utilized in behavioral selection in *Lymnaea*, we characterized the sensory pathway that conveys tactile information from the radula to the feeding network. Specifically, we identified a bilaterally-located pair of neurons in the buccal ganglia that both projected to the radula structure (Fig. 5a, b) and responded to brief tactile stimulus with robust activity. We confirmed that these neurons had a primary mechanosensory function by demonstrating that this radula tactile response persisted when chemical synaptic transmission was blocked (Supplementary Figure 5a). Importantly, sensory-driven spikes in these neurons, which we termed radula mechanosensory cells (RM), evoked large excitatory inputs on the command-like cell vTN, the key neuron-type that triggers feeding cycles in the presence of a stimulus during an appetitive bite (Crossley et al., 2016) (Fig. 5c). To test whether changes in hunger-state caused alterations in the response properties of RMs and in turn vTNs, we recorded these cells in fed or food-deprived animal preparations during radula stimulation. Strikingly, prior feeding state had no influence on the number of spikes elicited in RM cells, their amplitude or their inter-spike interval in evoked bursts (Fig. 5c, d, amplitude: fed, 37.5 ± 3.2 mV, food-deprived, 37.2 ± 2.1 mV, Mann Whitney test p = 0.88, inter-spike interval: fed, 0.06 ± 0.01 s, food-deprived, 0.055 ± 0.004 s, Mann Whitney test p = 0.96). Moreover, there was no effect on the amplitude of excitatory responses recorded on vTN (Fig. 5c-e). We also ruled out the possibility that modulation of dopamine signaling altered the sensory processing of the tactile stimulus by demonstrating that sulpiride treatment did not change either the RM response or the output to vTN (Supplementary Figure 5b-d). Taken together, these results suggest that the identified central switch mechanism is sufficient for behavioral selection in the absence of sensory retuning.

**Fig. 5.**
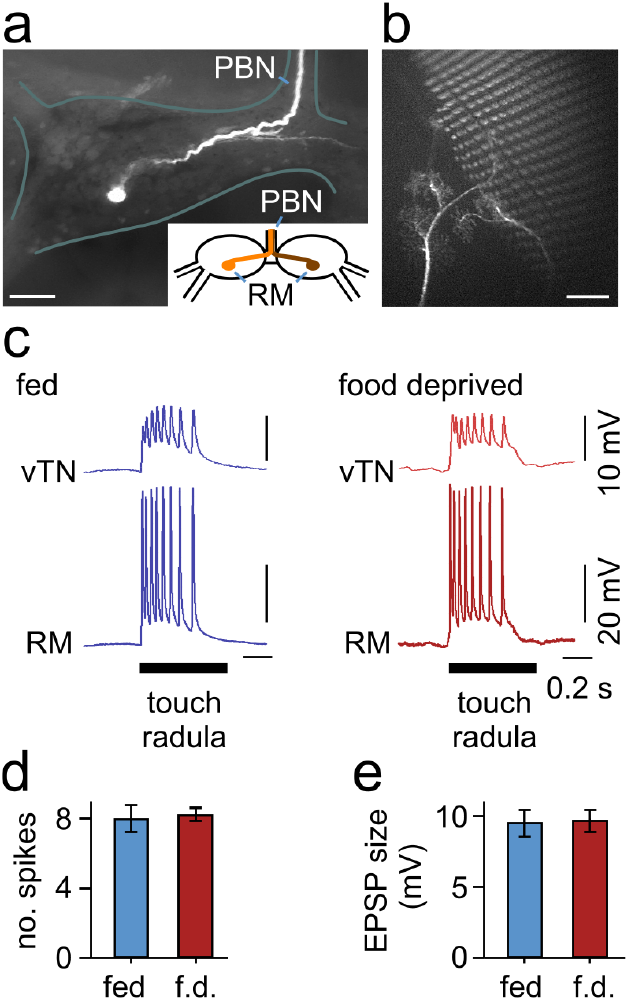
Hunger-state does not alter properties of mechanosensory neurons. (**a**) Morphology and schematic of radula mechanosensory neuron (RM) in buccal ganglia. A single projection leaves the ganglion via the post buccal nerve (PBN) towards the radula. Morphology confirmed in n = 4 preparations. Scale bar 100 μM. (**b**) Image of a single RM’s projection under the toothed radula. Scale bar 100 μM. (**c**) Representative trace of RM and vTN’s response to a tactile stimulus to the radula in fed (blue traces) and food-deprived (red traces) preparations. Touch induced RM spikes elicit 1:1 EPSPs on vTN. Black bar represents duration of tactile stimulus (0.5 s). (**d-e**) Statistical analysis of RM and vTN response to touch to the radula. There was no significant difference in the number of RM spikes in response to touch between fed (n = 8) and food-deprived (n = 8) preparations (Mann Whitney test, p > 0.05) or size of EPSP on vTN (unpaired t-test, p > 0.05). Error bars represent mean ± SEM.

### Central control allows generalization of behavioral choice to multiple cues

Given that the central switch mechanism we have characterized is functioning independently of sensory retuning (Fig. 5), we hypothesized that it might generalize to alternative input stimuli. To test this key idea, we used a behavioral forced-choice paradigm with a different sensory modality stimulus - in this case a chemical cue (amyl acetate, AA) applied to the mouth during an ingestive bite - thus ensuring no overlap in sensory processing pathways with our previous paradigm. Remarkably, in fed animals, the majority of the post-stimulus responses were classified as egestion, whereas in food-deprived animals, most were classified as ingestion (Fig. 6a, b). Thus, the hunger-state dependent alteration in the perceived value of a stimulus and subsequent behavioral selection generalizes to another sensory stimulus. Finally, we tested whether intervention with sulpiride was sufficient to bias fed animals towards ingestion in response to AA, replicating our findings with the tactile stimulus paradigm. We found that fed animals injected with sulpiride showed significantly more ingestive responses to AA than saline-injected animals (Fig. 6c, d). Therefore, the same drug treatment can alter the animal’s perceived value of stimuli of different modalities, providing final confirmation that PRN acts as a central control switch mechanism for motivational state-dependent behavioral selection.

**Fig. 6.**
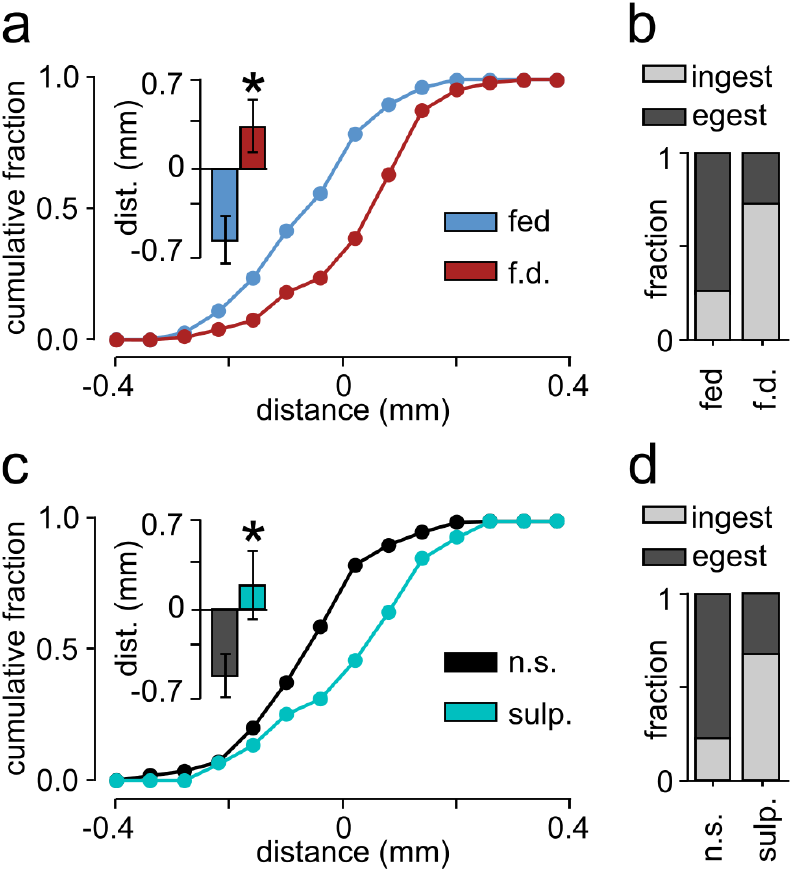
Hunger-state generalizes motor program selection to chemical stimulus. (**a**) Cumulative frequency plot of movements of the radula in response to amyl acetate (AA) in fed and food-deprived animals. There was a significant difference in the average radula movement in fed (−0.57 ± 0.19 mm, n = 20) vs food-deprived (0.34, ± 0.21 mm, n = 22) animals (unpaired t-test, p < 0.01). (**b**) Comparison of the fraction of ingestion and egestion responses to AA. There was a significant difference between fed and food-deprived animals (Fisher’s exact test, p < 0.0001). (**c**) Cumulative frequency plot of movements of the radula in response to AA in animals injected with saline or sulpiride. There was a significant difference in the average radula movement in saline (−0.52 ± 0.17 mm, n = 18) compared to sulpiride injected (0.19, ± 0.28 mm, n = 18) animals (unpaired t-test, p < 0.05). (**d**) Comparison of the fraction of ingestion and egestion responses to AA. There was a significant difference between saline and sulpiride injected animals (Fisher’s exact test, p < 0.0001).

## Discussion

An animal’s perception of key features of the environment can change dynamically based on its current internal state. In feeding behavior, this is a critical adaptive mechanism allowing the organism to make cost-benefit decisions about food value versus hunger state, even to the extent that a single stimulus can drive mutually exclusive responses (ingestion or egestion) according to motivation. How the nervous system computes and encodes information about input valence that can lead to the full reversal of behavioral action is not clear. Here we identify a central control switch in *Lymnaea* that addresses this demand, comprised of a dopaminergic interneuron type, PRN, which encodes hunger state, integrates sensory input and, through direct influence on the feeding network, biases an animal towards either ingestion or egestion in response to the same input. Our findings reveal an elegant solution for encoding a complex motivation-driven behavioral reversal with broad relevance for understanding how neural circuits can use parsimonious mechanisms to drive state-dependent peripheral decisions.

Previous examples of motivation-driven behavioral decision-making have revealed a key role for retuning of sensory pathways (Breunig et al., 2010; Ghosh et al., 2016; Su and Wang, 2014). In behavioral responsiveness paradigms in *Drosophila*, hunger increases feeding responses to sucrose (Inagaki et al., 2012; Marella et al., 2012) and movement towards certain odorants (Ko et al., 2015; Root et al., 2011), and decreases responsiveness to aversive stimuli (Inagaki et al., 2014; LeDue et al., 2016). Similar fine tuning of sensory processing in the leech has been implicated in the altered perception of stimuli in other state-dependent decisions (Gaudry and Kristan, 2009). Likewise, in comparable vertebrate paradigms, hunger state modulates sensory neuron firing properties in response to a stimulus (Breunig et al., 2010; Negroni et al., 2012; Nikonov et al., 2017). In all cases, for these relatively simple decisions where the animal is changing its responsiveness as a means to drive graded increases or decreases in behavior, sensory retuning is apparently an effective solution.

However, such examples contrast with the more complex decision-behavior task we examined in the present study. Here, an animal must switch entirely between two functionally-opposed behavioral outputs suggesting that fundamental differences in the neurobiological mechanisms might be required. Consistent with this, we experimentally ruled out the possibility that neurons involved in the detection of the tactile stimulus alter either their firing properties or their synaptic connections with command-like interneurons in response to different hunger states. Utilizing an isolated CNS preparation we demonstrate that hunger state persists in the complete absence of direct sensory activation, and biases the configuration of the network. Extending this idea further we show that, remarkably, this central control switch can also serve to generalize behavioral selection to alternative stimulus modalities. In mice, manipulating the activity in a subset of neurons in the lateral hypothalamus can alter the animal’s consummatory responses (Jennings et al., 2015; Nieh et al., 2015), even to the extent of promoting the ingestion of stimuli of non-calorific content or non-food stimuli (Burnett et al., 2016; Navarro et al., 2016). Although not hunger-state dependent, these studies demonstrate that, similar to *Lymnaea*, central mechanisms can be utilized to generalize behavioral selection to multiple sensory stimuli. In view of *Lymnaea’s* complex and varied natural sensory environment, we argue that this provides an efficient mechanism for an animal to set its general level of feeding readiness regardless of the specific stimuli that it encounters.

Our findings suggest a level of network degeneracy (Beverly et al., 2011; Gutierrez et al., 2013), where distinct circuit mechanisms can generate the same motor output in a context-dependent manner. The master-switch for hunger-state dependent ingestion/egestion behavioral selection is specific to this role whereas a separate pathway is necessary for esophageal driven egestion. Indeed, we found that this purely egestive stimulus was not modulated by the animal’s hunger state. This kind of degeneracy, similar to examples in other vertebrate and invertebrate feeding circuits (Betley et al., 2013; Cropper et al., 2016), provides both flexibility and hard-wiring of behavior, allowing adaptive responses to a stimulus to be based on the animal’s internal state but additionally deprioritizing the hunger induced bias towards ingestion in response to certain potentially harmful stimuli. Similar mechanisms are utilized in mammals, where hunger state can alter the perception of inflammatory pain while transient nociceptive pain remains unaffected (Alhadeff et al., 2018). As in *Lymnaea*, this mechanism allows for behavioral adaptation during times of food-deprivation, but not at the expense of exposing the animal to potentially life-threatening encounters.

In most systems investigated so far, dopamine has been found to play a role in the processing of a variety of reward-related behaviors (Harris et al., 2012; Kemenes et al., 2011; Marella et al., 2012; Schultz, 2016; Stauffer et al., 2016; Waddell, 2013). It is notable therefore, that the central-switch neurons, the PRNs, which drive egestion in fed animals, are dopaminergic. Dopaminergic action has been shown to cause network reconfiguration in other systems (Marder, 2012; Neveu et al., 2017) as well as playing an important role in shaping behavioral output in a hunger-dependent manner (Inagaki et al., 2012; Marella et al., 2012). Interestingly, in these examples, dopamine causes an increase in the positive valuation of a stimulus, whereas in the *Lymnaea* forced-choice paradigm it seems to lead to a more negative valuation. However, based on the opponent process theory of motivation (Solomon and Corbit, 1974), successful avoidance of a potentially dangerous or unpleasant situation (e.g., one that results in the ingestion of a non-palatable object) may be a reward in its own right (Schultz, 2015). This notion is supported by our finding in *Lymnaea* that dopamine plays an important role in making the decision to reject a potentially harmful stimulus.

Collectively, our findings reveal a fundamental mechanism for changing the perceived value of potential food stimuli in the environment, weighted according to the animal’s motivation to feed. A central control mechanism allows the circuit to lower its threshold for promoting ingestion behavior when the animal is hungry, even generalizing across different input modalities, and thus raising the likelihood of successful foraging but at the risk of consuming inedible and potentially harmful foods. In broader behavioural terms, this mechanism would therefore serve to allow an animal to use a riskier feeding strategy when increased hunger state demands it.

## Acknowledgements

We thank Michael Schofield and Paul R. Benjamin for help with the dopamine staining. This work was funded by the Biology and Biotechnology Research Council (BBSRC/BB/H009906/1 and BBSRC/BB/P00766X/1) to G.K. and M.C. K.S. was supported by funding from BBSRC/BB/K019015/1.

## Author Contributions

M.C., K.S. and G.K. conceived and designed the experiments. M.C. performed the experiments. M.C. and K.S analyzed the data and made figures. M.C., K.S. and G.K. wrote the paper. G.K. acquired the funding and was responsible for resources.

## Materials & Correspondence

All correspondence and material requests should be addressed to

## Methods

### Animal maintenance

All experiments were performed on adult (3-4 months old) *Lymnaea stagnalis*. Animals were kept in groups in large holding tanks containing Cu^2+^ free water at 20°C on a 12:12h light-dark regime. The animals were fed lettuce three times a week and a vegetable based fish food (Tetra-Phyll; TETRA Werke, Melle, Germany) twice a week. Animals were transferred to smaller holding tanks prior to experimenting. Animals were either fed *ad libitum* or food-deprived for four days prior to either behavioral or electrophysiological experiments.

### Preparations and electrophysiological methods

*In vitro* experiments were carried out using a buccal mass-CNS preparation(Rose and Benjamin, 1979) or a radula-CNS preparation (Crossley et al., 2016). The buccal mass-CNS preparations was used to record from muscles extracellularly and neurons intracellularly within the CNS. Experiments in Fig. 1c, d were carried out on a preparation that consisted of the entire buccal mass attached to the CNS via the left and right lateral and ventral buccal nerves (LBN and VBN respectively), enabling the co-recording of the SLRT and AJM muscles and intracellular recording of command-like interneuron CV1a (McCrohan, 1984). A small region of the anterior esophagus was kept attached to the CNS via the dorsal buccal nerves (DBN). A modified version of this preparation was used in Fig. 1h-l and Fig. 2-4 which consisted of the buccal mass connected to the CNS via the left LBN and VBN only. During these recordings, the AJM was the only muscle recorded from. This preparation provided greater stability for performing intracellular recordings from moto- and interneurons within the buccal ganglia whilst still providing a readout of the retraction phase. The radula-CNS preparation was used to identify mechanosensory neurons and test whether motivational state altered their response to tactile stimulation of the radula. A 0.5 s tactile stimulus was applied to the radula via a mechanical probe controlled by the CED ADC board (micro 1401 Mk II interface - Cambridge Electronic Design, Cambridge, UK). A lip-CNS preparation was used in Supplementary Figure 1a to confirm CV1a’s responsiveness to appetitive stimuli to the lips. The preparation used is described in detail in (Kemenes et al., 2001). Saline containing 0.67% sucrose was applied to the lips whilst recording intracellularly from CV1a. A modified version of this preparation was used in Fig. 1c where the buccal mass was left attached to the CNS similar to that used in Fig. 1c, d. This allowed the recording of muscles whilst applying appetitive stimuli to the lips. Preparations were perfused with normal saline containing 50 mM NaCl, 1.6 mM KCl, 2 mM MgCl2, 3.5 mM CaCl2, 10 mM HEPES buffer in water. Intracellular recordings were made using sharp electrodes (10-40 MΩ) filled with 4 M potassium acetate. NL 102 (Digitimer Ltd) and Axoclamp 2B (Axon Instrument, Molecular Device) amplifiers were used and data acquired using a micro 1401 Mk II interface and analyzed using Spike2 software (Cambridge Electronic Design, Cambridge, UK). Muscles were recorded using a glass suction electrode. Signals were amplified using an NL104 (20 k gain) (Digitimer Ltd) and were low pass (50 Hz) and notch (50 Hz) filtered using NL125/126 filters (Digitimer Ltd) before they were digitized at a sampling rate of 2 kHz using a micro 1401 Mk II interface (Cambridge Electronic Design, Cambridge, UK).

### Neurons and muscles recorded

The anterior jugalis muscle is a large thick muscle which is involved in the retraction of the buccal mass and the radula/odontophore complex (Carriker, 1946; Rose and Benjamin, 1979; Staras et al., 1998). Electromyography (EMG) recordings were obtained from the anterior region of this muscle. The majority of activity recorded occurs during the retraction phase (Rose and Benjamin, 1979). The supralateral radula tensor muscles are the largest of the tensor muscles and the bulkiest in the odontophore and have been previously reported to be involved in the retraction phase of a cycle (Carriker, 1946; Rose and Benjamin, 1979). EMG recordings were performed on the SLRT on the dorso-lateral edges of the odontophore. Command-like interneuron CV1a is located in the cerebral ganglia and was identified by its electrical properties, characteristic location and its ability to drive fictive feeding cycles when artificially depolarized to fire spikes (McCrohan, 1984; McCrohan and Kyriakides, 1989). The N2v neuron is a CPG interneuron located on the ventral surface. It can be identified by its characteristic plateau during the retraction phase of a cycle. Artificial activation of an N2v causes wide spread retraction phase activity in many buccal neurons (Brierley et al., 1997a; Brierley et al., 1997b), and activity on the AJM. B11 is a newly identified SLRT motoneuron located on the ventral surface of the buccal ganglia. Spikes in B11 caused contraction of the SLRT muscle. Touch to the esophagus initiated a barrage of EPSPs on B11 which caused it to spike in the protraction phase of the initiated cycle. PRN is a newly identified buccal-cerebral interneuron which can be identified via its ability to drive fictive feeding cycles, its activation during cycles initiated by touch to the esophagus and via its 1:1 excitatory connection with B11. RM is a newly identified mechanosensory neuron in the buccal ganglia. Touch to the radula causes somatic spikes in RM which rise from baseline and persist in a saline containing zero Ca^2+^ and ethane glycol-bis (β- aminoethylether)-N, N, N’,N’-tetracetic acid (EGTA), blocking chemical synaptic transmission. The saline contained: 35.0 mM NaCl, 1.6 mM KCl, 18.0 mM MgCl2, 2.0 mM EGTA, and 10 mM HEPES buffer in water. vTN was identified due to its location and white color and response to tactile stimulation of the radula (Crossley et al., 2016).

### In vitro classification of cycles

Artificial activation of CV1a was used to drive fictive ingestion cycles. CV1a is a command-like interneuron that is activated by appetitive sensory stimuli to the lips which drive ingestion behavior *in vivo* (Kemenes et al., 2001; Kemenes et al., 2002; Whelan and McCrohan, 1996) and see Supplementary Figure 2a). To drive ingestion cycles *in vitro*, depolarizing current was injected into a single CV1a to elicit spiking until cycles were initiated. These cycles were indistinguishable from those generated by application of appetitive stimuli to the lips (sucrose). To elicit egestion *in vitro*, a 1 s tactile stimulus was applied to the region of the esophagus proximal to the point of entry of DBN from the buccal ganglia. The esophagus contains fibers from mechanosensory neurons which provide aversive cues to the feeding system (Elliott and Benjamin, 1989). The tactile stimulus was applied to the esophagus using a mechanical probe controlled by a TTL pulse from the micro 1401 Mk II. Large high frequency AJM activity only occurs in the retraction phase (Benjamin and Rose, 1979; Rose and Benjamin, 1979; Staras et al., 1998), therefore we used this as a constant phase of activity with which to measure the onset of the retraction phase in all preparations. AJM activity was plotted in 0.2 s bins to aid in identification of the onset of the burst, signaling retraction phase initiation. Activity on the SLRT muscle was measured with respect to the onset of the AJM burst. To analyze the relative activity of SLRT, it was measured 5 s before and 5 s after the AJM burst onset. The number of SLRT events before AJM onset was subtracted from the number of SLRT events after AJM onset and then divided by the total number of events in the 10 s period to gain a normalized difference score. Using this normalized difference score, a positive score represents more activity occurring after AJM onset, and a negative score represents more activity occurring before AJM onset. Based on analysis of those cycles driven by either CV1a and touch to the esophagus, cycles were classified as fictive ingestion cycles if a greater proportion of activity occurred after AJM onset and fictive egestion a greater proportion of activity occurred before the AJM burst onset. The same form of analysis and criteria were used for classifying cycles in which B11 was co-recorded with AJM. B11 spike activity was also plotted as heat plots in Fig. 2-4 using MatLab software. B11 spikes were binned in 0.2 s bins. Data was organized from cycles with most to least activity prior to AJM onset. To compare the effects of satiety on fictive feeding cycles *in vitro*, the first 10 spontaneous cycles were analyzed from 12 fed preparations and 12 food-deprived preparations. To test the effects of sulpiride on cycles in fed preparations, the first 10 cycles were recorded in each preparation and then 10^-4^ M sulpiride in normal saline was perfused on the preparation. The first 10 cycles which occurred after 10 min of perfusion were analyzed. PRN spike frequency in fed or food-deprived preparations was recorded in the first 10 spontaneous cycles which occurred in each preparation. PRN spike frequency was analyzed for 5 s before the onset of the retraction phase and plotted in 0.25 s bins.

### Behavioral paradigms

#### Behavioral paradigm 1

*Lymnaea*’s behavior was observed by placing them in a custom built behavioral chamber filled with Cu^2+^ free water. The chamber held the animal on the surface of the water allowing for the application of sensory stimuli to the mouth of the snail whilst being able to fully observe movements of the feeding structures. All behavioral experiments were videoed and analyzed using ImageJ. Animals were left to acclimatize for 10 min prior to testing. An ingestion bite was triggered by brief application of an appetitive stimulus (lettuce) to the lips of the animal, eliciting a bite response in all animals tested, regardless of hunger-state (148 animals). Upon opening of the mouth either a tactile probe was placed inside or 50 μL 0.008% amyl acetate (AA) was applied to the mouth/lips of the animal. The tactile probe consisted of a 1-ml syringe whose tip had been heated and pulled into a fine point. In both conditions, a radula motor-program was induced in response to the stimulus. All animals were videoed and these responses later analyzed (see below section for details of analysis).

#### Behavioral paradigm 2

To test whether tactile stimulation of the esophagus elicited ingestion or egestion, an incision was made under the mantle cavity to expose a region of the esophagus, allowing for the mechanical stimulation of the structure with a pair or forceps. Esophageal stimulation was sufficient to elicit an egestion bite even when presented during a period of quiescence. The elicited bite was videoed and analyzed (see below).

#### Behavioral paradigm 3

To determine the effects of drug injection on animal’s locomotion we tracked animals in a novel environment for 30 mins. Briefly, animals were placed in a 14 cm-diameter petri dish filled with 100 ml Cu^2+^ free water. Recording started as soon as they were placed in the arena so as to monitor their initial behavior. Animals were recorded at 1 frame/s for 30 min. Videos were analyzed using idTracker (Perez-Escudero et al., 2014) and total distance traversed compared between groups.

#### Behavioral paradigm 4

The effects of drug treatment on the animal’s food searching behavior in a novel environment were tested by counting the number of appetitive bites during the first 10 min from being placed in a petri dish filled with 100 ml Cu^2+^ free water (as in(Kemenes and Benjamin, 1994).

#### Behavioral paradigm 5

The effects of drug treatment on the animal’s responsiveness to sucrose was tested by placing the animal in a petri dish of 90 ml Cu^2+^ free water. Animals were allowed to acclimatize for 10 min and then 5 ml water was added to the dish. The number of bites performed were counted for 2 min then 5 ml sucrose (0.33% final solution) was added and the number of bites performed counted for 2 min. A feeding score was obtained by subtracting the number of bites in response to water from the number performed in response to sucrose.

### Analysis of behavior and characterization of ingestion and egestion in vivo

Biting behavior was videoed at 33 frames/s. The direction of movement of the radula and underlying odontophore were measured during the response to either the tactile probe or AA. The video was first rotated so that the animal was aligned with their head to foot in an anterior to posterior direction. From the first frame where the radula was visible, it was tracked using ImageJ software in the y-axis of movement for 10 frames. The last frame was subtracted from the first frame to give a positive or negative direction of movement. A positive score therefore represented a majority anterior movement of the radula whereas a negative score represented a majority posterior movement. We further measured each of the 10 points the radula was tracked and each point was subtracted from the subsequent point and cumulative plots of radula movements were generated. We set the criteria to define whether a response was ingestion or egestion based on whether the difference in movement was positive (ingestion) or negative (egestion).

### Iontophoretic dye filling of neurons

Neurons were filled with a fluorescent dye allowing the morphology of the cell to be determined. Microelectrodes were filled with either 5(6)-carboxyfluorescein (5-CF) or AlexaFluor 568 (Molecular Probes). Cells were filled iontophoretically using a pulse generator which applied regular interval negative square current pulses into the neuron for >30 min. The preparation was then left overnight at 4°C. Images of the neurons were taken using a digital camera (Andor Ixon EMCCD) mounted on a Leica stereomicroscope.

### Dopamine immunohistochemistry whole mount staining

CNSs were dissected out in normal saline with 1% sodium metabisulfite (MBS) and neurons identified and filled with AlexaFluor 568 as above. The preparations (n = 4) were then incubated in 0.25% Protease XIV (Sigma, Aldrich) for 5 mins in saline/MBS at room temperature and then washed with saline/MBS. Next, the preparations were fixed for 1 h in fixative solution (5% glutaraldehyde in 0.1 M sodium cacodylate buffer (pH 7.4)) at room temperature and then washed in Tris/MBS buffer (0.1 M Tris base, 0.15 M NaCl) and reduced with 1% sodium borohydride in Tris/MBS for 10 min and washed x3 with Tris/MBS. They were then incubated in 0.1 M phosphate buffer with 4% Triton-X (PBT) plus 1% MBS for 4 h. Blocking was performed with 1% bovine serum albumin (BSA) in 0.25% PBT plus 1% MBS overnight at 4°C. The blocking reagent was removed and the preparations were then incubated with fresh blocking reagent as above but with rabbit anti-dopamine (AbCam) at 1/1000 and incubated for 72 h at 4°C. Three washes with 0.25% PBT followed. The preparations were blocked with 1% normal goat serum in 0.25% PBT plus 1% MBS for 4 h at room temperature. The blocking reagent was removed and the preparations were incubated with fresh blocking reagent as above plus goat anti-rabbit Alexa 488 (Invitrogen) at 1/100 for 48 h at 4°C. Three washes with PBS followed. Preparations were then mounted in glycerol mountant on a cavity slide to be imaged.

### D2 receptor blocker application in vitro and in vivo

To test for the effect of sulpiride (±) on the PRN→B11 connection, preparations were bathed in high divalent (HiDi) saline which increases the threshold for action potentials, acting to reduce polysynaptic connections. HiDi saline was composed of 35.0 mM NaCl, 2 mM KCl, 8.0 mM MgCl2, 14.0 mM CaCl2, and 10 mM HEPES buffer in water. Baseline EPSP size was acquired and then 10^-4^ M sulpiride (±) (Sigma, Aldrich) in HiDi saline was perfused into the bath for 10 min and EPSP size recorded again. To test the behavioral effects of sulpiride, animals were injected with 100 μL of 10^-3^ M sulpiride (±) in normal saline. The injected concentration of the drug was diluted ~10-fold by the body fluids of the animal(Ford et al., 2015). Control animals were injected with 100 μL normal saline alone. Animals were left for 2 h before behavioral tests were carried out.

### Quantification and statistical analysis

Data was analyzed using GraphPad Prism 5 (GraphPad Software) and expressed as mean ± SEM. Each ‘n’ represents an individual preparation unless stated otherwise in the text. Normality was tested using the D’Agostino and Pearson omnibus normality test. Where data was shown to be normally distributed, two-group statistical comparisons were performed using two tailed t-test statistics (either paired or unpaired as stated in the text). Non-normally distributed data was analyzed using a Mann-Whitney U-test or Wilcoxon signed rank test. The comparisons between the percentage of bites/cycles classified as ingestion or egestion was compared using a Fisher’s exact test. The significance level was set at p < 0.05.

### Data availability statement

The data that support the findings of this study are available from the corresponding author upon reasonable request.

